# Sterile insect technique reduces cabbage maggot (Diptera: Anthomyiidae) infestation in root crucifers in Canada

**DOI:** 10.1101/2025.05.09.653057

**Authors:** Anne-Marie Fortier, Allen Bush-Beaupré, Jade Savage

## Abstract

The cabbage maggot (*Delia radicum* Linnaeus) is a major pest of brassica vegetables in Canada that has traditionally been managed with soil-applied insecticides. However, recent regulatory restrictions on key products such as chlorpyrifos have created a pressing need for alternative solutions. This study evaluates the sterile insect technique (SIT) as a control method for the cabbage maggot in root crucifers. Large-scale field trials conducted from 2019 to 2022 in Quebec (Canada) demonstrated significant reductions in *D. radicum* infestations in radish and daikon crops. Quality control measures confirmed the effectiveness of sterilization, with minimal impact on male performance. The results suggest that the SIT is a promising, environmentally friendly alternative to chemical control for cabbage maggot management. The study further highlights the importance of optimizing release strategies and improving predictive models to guide deployment. Overall, the SIT offers growers a viable option to reduce reliance on insecticides while maintaining crop health and yield.

---

The *Brassicaceae* family includes approximately 3,700 species, many of which are of agronomic importance as vegetables, condiments, fodder and oil crops (Beilstein et al. 2006). Canada produces more than 15,000 hectares of the main brassica vegetables (cabbage, cauliflower, broccoli, rutabaga and radish), with the province of Québec accounting for about 50 % of the national production (Agriculture and Agri-Food Canada 2024). The cabbage maggot (CM), *Delia radicum* (Linnaeus) (Diptera: Anthomyiidae), is a major threat to brassica crops in the temperate northern hemisphere (Finch 1989, Walgenbach et al. 1993, Dixon et al. 2014). It overwinters as diapausing pupa in the soil (Turnock et al. 1985, Johnsen et al. 1997) and in spring, adult flies emerge, mate, and females lay eggs singly or in small clusters at the base of the host plant (Matthewman and Harcourt 1971, Dosdall et al. 1994). Plant damage occurs when hatched larvae burrow into the soil to feed on root hairs and subsequently enter the taproot (Richard and Boivin 1994). Feeding damage can facilitate the entry of pathogens into injured roots, and often result in slow growth, yellowing, stunting and eventually plant death (Santolamazza-Carbone et al. 2017). When severe infestations occur, yield losses can reach up to 100%, especially if young plants are affected (Ferry et al. 2009). Root crucifers like radish and rutabaga are particularly vulnerable to CM, since the damage directly affect product quality and shelf life. Annual economic losses due to CM infestations in Northern America are estimated to be $100 million (Wang et al. 2016).

Due to their belowground immature life stages, CM can be difficult to control. Growers have typically applied granular and/or drenched insecticides to the soil but the number of available insecticides for CM control in Canada and elsewhere has diminished greatly over the past decades. Until recently, chlorpyrifos was the most widely used insecticide and considered the most effective, although cases of resistance have been confirmed in rutabaga production areas (Van Herk et al. 2016). However, this organophosphate is deleterious to the health of users and consumers, having been notably associated with autism and IQ deficits in children (Rauh et al. 2012, Bellanger et al. 2015) and identified as a major contaminant of surface water in vegetable production areas (Hunt et al. 2003, Giroux and Fortin 2010). Due to environmental and human health considerations, all products containing chlorpyrifos in Canada were deregistered by the Pest Management Regulatory Agency (PMRA) (Health Canada, REV2021-04) with last date of use set to December 2023. The only active ingredients currently available to control the CM are cyantraniliprole (Verimark®) and spinosad (Entrust®, Success®), but the number of applications allowed in a season is limited, and they are also used to control other important insect pests of these crops. Alternative control methods for the CM such as the deployment of trap crops, predators, and microbial agents exist but are often considered impractical or provide insufficient or inconsistent control, and few have been implemented commercially (Collier et al. 2020). The most widely non-insecticidal control method used is the deployment of physical barriers (net covers), but they generally must be removed before the end of egg-laying risks to avoid adverse effects (increase in temperature and humidity, loss of light) and allow weeds or pathogens management. It is therefore essential to develop or optimize alternative control methods for this pest that will minimize potential negative impacts on non-target organisms, human health and the environment.

The sterile insect technique (SIT) is an environmentally friendly pest management tool that has been successfully applied worldwide to manage agricultural pests such as the codling moth *Cydia pomonella*, the Mediterranean fruit fly *Ceratitis capitata*, and the onion maggot *Delia antiqua* (Klassen and Curtis 2005, Dyck et al. 2021). This method typically involves the mass-rearing of the target pest species, sterilization using gamma or x-rays radiation, and sustained release in the target area in sufficient numbers to achieve appropriate sterile to wild insect ratios. Wild females mated with sterile males then lay sterile eggs, thereby resulting in a rapid decline in pest populations. Since 2005, the SIT has been developed and implemented for onion maggot control in the province of Québec (Fournier and Brodeur 2012), resulting in the gradual abandonment of chlorpyrifos incorporation at sowing for most SIT users before it was deregistered (Fortier et al. 2024). The larger-scale releases for sterile onion maggot began in 2011 and resulted in a significant decrease of watershed contamination by chlorpyrifos (Giroux 2017).

Previous small-scale field-based studies have provided encouraging results regarding the potential feasibility of using the SIT as a tool for CM control (Fournier and Fortier 2014; Fortier et al. 2017). The monitoring of CM populations on a daikon (Chinese radish) farm in the Montérégie region (Quebec, Canada) where sterile CM were released between 2015 and 2019 showed an important decrease in the density of wild populations and significant reductions in release rates of sterile flies over the monitoring period (Fortier, unpublished data). Building on these preliminary results, the goal of the present study was to evaluate the performance of the SIT as an alternative method to control the cabbage maggot and reduce infestation rates in large-scale field trials conducted in commercial conditions in radish and daikon crops over a four-year period (2019-2022). In addition, since the foundation of a successful SIT program lies in effective sterilization with minimal detrimental effects on male performance (Robinson 2005, Bakri et al. 2021), quality control results on the impacts of irradiation on emergence, sterility, and mating competitiveness of CM adults produced between 2015 and 2022 are also presented.

## Materials and methods

### Study area

The efficiency of sterile fly releases to control *D. radicum* was evaluated in 22 daikon and five radish fields across four commercial farms located in West Montérégie, QC, Canada between 2019 and 2022 (Table 1) (Farm 1: 45.20N, 73.36W; Farm 2: 45.13N, 73.63W; Farm 3: 45.15N, 73.59W; Farm 4: 45.17N, 73.56W). Farm 1 (daikon) had been exposed to sterile CM releases between 2015-2019 while farms 2-4 (radish) had no prior exposure. All radish fields and 10 of the 22 daikon fields were compared with untreated control fields located on the same farm, sown at the same time with the same variety, and separated by a buffer zone of at least 300 meters. All fields were characterized by organic soils (muck soil) and under conventional management.

**Table 1.**
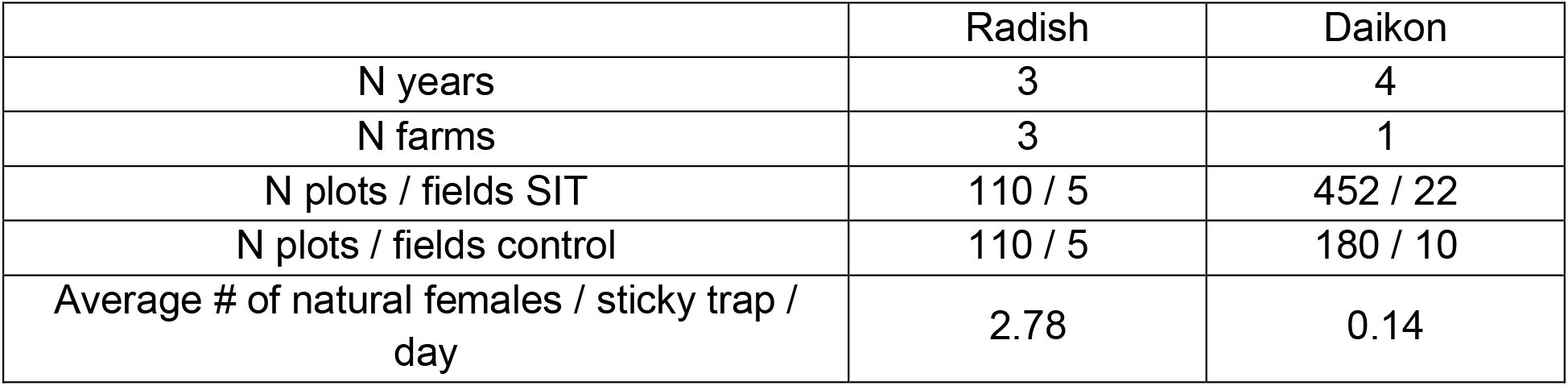
Sample sizes for the evaluation of sterile fly release efficiency for the control of the cabbage maggot in daikon and radish crops.

|  | Radish | Daikon |
| --- | --- | --- |
| N years | 3 | 4 |
| N farms | 3 | 1 |
| N plots / fields SIT | 110 / 5 | 452 / 22 |
| N plots / fields control | 110 / 5 | 180 / 10 |
| Average # of natural females / sticky trap / day | 2.78 | 0.14 |

### Mass production

Our laboratory colony of *D. radicum* was first established in 2012 using flies emerging from pupae collected in naturally infested radish and daikon plants in West Montérégie; since then, the colony has been refreshed yearly with wild specimens from the same area. All sterile CM released in the context of this work were produced at Phytodata mass rearing facility (Sherrington, Quebec, Canada) using procedures adapted from Vereecke and Hertveldt (1971) and Van Keymulen et al. (1981). The adults were kept in 46 × 46 × 40 cm cages covered with nylon mesh walls and supplied with osmosed water and a 10:10:2:1 mixture of powdered milk, powdered sugar, brewer’s yeast and soya flour. Pieces of rutabaga, *Brassica napus* variety *napobrassica* L. (Brassicaceae) served as oviposition sites. Eggs were collected three times per week and inoculated on 25 × 51 × 6 cm plastic trays containing 4 kg of rutabaga (0,15 g/kg). Larvae were reared during 25 days at 20.5 ± 1.0 °C and about 70 ± 2 % relative humidity and a 16L:8D photoperiod. Harvested pupae were kept at 4°C until irradiation. From 2019 to 2021, the pupae were gamma-irradiated with Cobalt-60 by Nordion (Laval, QC, Canada) at 35 Gy. In 2022, pupae were irradiated at Phytodata with the same dose, using an RS 1800Q X-ray machine (Rad Source Technologies, Georgia, USA). Irradiated pupae were colored with DayGLo® pink dye ECO-11 (Debro, ON, Canada) to facilitate field identification of emerged adults, and placed in an emergence box with a source of water and food. Emerged flies were fed *ad libidum* for 3 days, before being packaged and released.

### Releases

The sterile flies were manually released on a weekly basis over a period of five to ten weeks along the borders surrounding each of the 27 fields, with weekly quantities adjusted based on the activity and abundance patterns of wild populations at rates ranging from 9 500 to 85 000 flies per hectare depending on pest pressure and time of season, with higher rates early in the season, corresponding to the first CM generation (Table S1). For daikon, release rates were globally lower given the reduced populations through the previous four years of releases. For radish, the participating farms were using sterile CM flies for the first time and wild population densities were initially unknown, so a higher release rate was targeted. Nonetheless, yearly release rates were ultimately determined by the maximum amount that growers were willing to pay. No insecticide sprays were applied to control the CM in the experimental fields for the duration of the study.

### Field monitoring and damage assessments

All SIT and control fields were monitored for adult catches and maggot damage between first release and harvest time. Three yellow 10 × 16 cm sticky cards (Distributions Solida, QC, Canada) with the food attractant CSALOMON for *Delia radicum* (Plant Protection Institute, Hungary) were installed approximately 100 meters apart in field borders and collected twice a week for examination, specimen counts, and determination of the sterile/wild fly ratio. At harvest, 10 consecutive plants were evaluated in 20 randomly selected plots in each field to estimate the cabbage maggot damage incidence. Larvae were collected and identified using Savage et al. (2016).

### Quality control

Between 2015 and 2022, adult emergence, sterility of irradiated males and females along with male mating competitiveness were evaluated for each weekly release. Four different combinations of male and female pairs (10 males and 10 females per combination) were formed: 1) non-irradiated males and females (control); 2) non-irradiated males and irradiated females (control of female sterility); 3) irradiated males and non-irradiated females (control of male sterility) and 4) 10 non-irradiated females combined with 5 irradiated males and 5 non-irradiated males (male competitiveness). For each combination, eggs were collected every 2-3 days over a 14 days period and incubated 5-7 days on petri dishes with moist filter paper to evaluate their hatch rates. A sample of at least 300 irradiated and non-irradiated pupae was kept for each weekly release to estimate the percentage of adult emergence.

### Statistical analyses

All generalized linear hierarchical models (GLHMs) described below were fit with glmmTMB (v.ref), model diagnostics evaluated with DHARMa (v.ref), marginal predictions and contrasts calculated with marginaleffects (v.ref) and plots produced with ggplot2 (v.ref) packages within the R environment (v.ref).

### Quality control

We fit separate GLHMs for adult emergence and egg hatchability. Both adult emergence and egg hatchability data had the same hierarchical structure. As treatment replicates originated from a sample from the same release event for each year, we included a random intercept of release ID nested within year to control for the non-independent measures of the treatment for each release. After model fitting, the contrast between the control and treatments were computed by marginalizing across the hierarchical structure of the models. The data on the proportion of hatched eggs out of the total number of eggs laid by females were modeled assuming a betabinomial error distribution with logit link function and had treatment (control vs sterile males & competition) as predictor. The data on adult emergence were modeled using a beta error distribution with logit link function and had treatment (control vs sterile) as predictor.

### Field trials

The data for radish and daikon were analysed with separate models which differed only with regards to the specification of the hierarchical structure of the data. Both models assumed a betabinomial error distribution with logit link function and had treatment (control vs SIT) as main predictor and the average number of natural females per sticky trap per day for the season used as a control variable for the magnitude of *D. radicum* infestation pressure for each field. As each radish field was evaluated for damage only once per farm, we used field ID as the sole random intercept in that model and thus represents the variance across fields, growers and years. In daikon, however, some fields were evaluated more than once across years and thus, a random intercept per field nested within year was used for that model. After model fitting, the contrasts between the control and SIT treatments were computed by setting the average number of natural females per sticky trap per day for the season to its mean and marginalizing across the hierarchical structure of the model. In other words, the predictions and contrasts reported are for an average field with an average *D. radicum* infestation potential. The average effect of the average number of natural females per sticky trap per day for the season was computed by calculating the average predictions for both treatments.

## Results

### Quality control

Model predictions, contrasts and outputs are reported in Figure 1 and Table 2 for both measures of quality control. For adult emergence from puparia, the predicted mean proportion of emerged adults [95% confidence intervals] was 80.1% [72.7, 85.8] for the control and 75.1% [66.7, 81.9] for the sterile treatment (Figure 1A). The percentage-point contrast between treatments (sterile vs control) was -4.99% [-7.28, -2.70] (p-value < 0.001; Figure 1B).

**Table 2.** Output of statistical analyses (generalized linear hierarchical models) for the effect of male group composition (sterile males, sterile males + non-sterile males (Competition), and non-sterile males (Control)) on female egg hatchability and effect of sterilization vs control on adult emergence from puparia of *D. radicum*. Output reports estimates, corresponding 95% confidence intervals (CI), *Z* statistic and *p*-values. A betabinomial error distribution with a logit link function was used for both GLHMs. σ^2^ represents the variance of the random effects (in subscript next to value). Note that the estimates are on the logit scale.

|  | Egg Hatchability |  |  |  | Adult Emergence |  |  |  |
| --- | --- | --- | --- | --- | --- | --- | --- | --- |
| *Predictors* | *Estimate* | *CI (95%)* | *Z* | *p* | *Estimate* | *CI (95%)* | *Z* | *p* |
| (Intercept) | 1.22 | 0.91 – 1.52 | 7.76 | **<0.001** | 1.39 | 0.98 – 1.80 | 6.61 | **<0.001** |
| Competition vs Control | -1.47 | -1.72 – -1.22 | -11.55 | **<0.001** |  |  |  |  |
| Sterile vs Control | -4.58 | -4.96 – -4.21 | -23.95 | **<0.001** |  |  |  |  |
| Sterile vs Control |  |  |  |  | -0.29 | -0.40 – -0.17 | -4.84 | **<0.001** |
| **Random Effects** |  |  |  |  |  |  |  |  |
| $\sigma^2$ | 0.06 release:year | | | | 0.42 release:year | | | |
|  | 0.09 year |  |  |  | 0.31 year |  |  |  |
| Observations | 395 |  |  |  | 314 |  |  |  |
| Marginal R<sup>2</sup> / Conditional R<sup>2</sup> | 0.882 / 0.923 |  |  |  | 0.027 / 0.983 |  |  |  |

**Figure 1.**
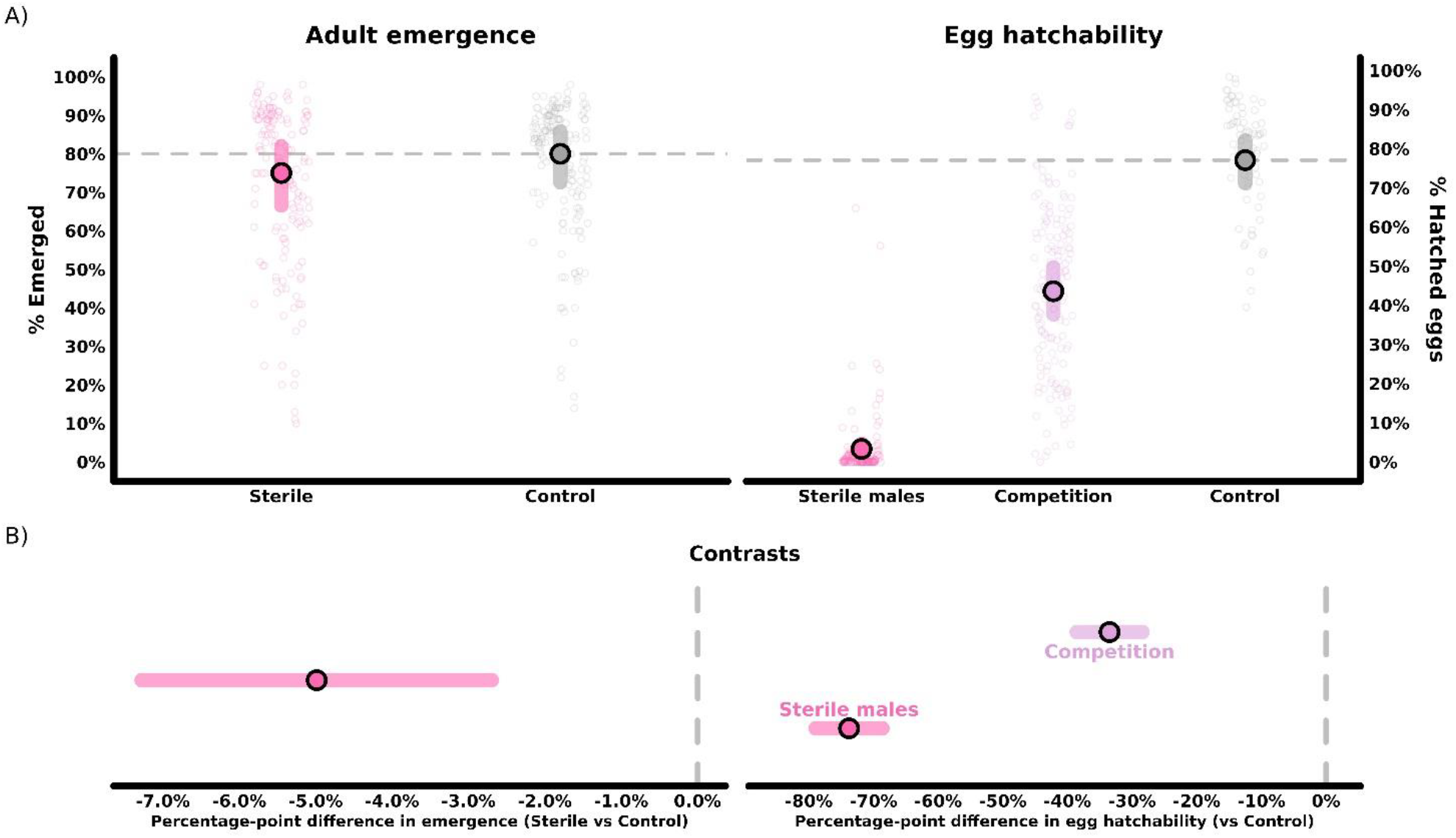
Marginal predictions of percent D. radicum adult emergence and eggs hatched for the sterile & control (N = 157) treatments (adult emergence) and sterile males (N = 163), competition (N = 159) & control treatments (N = 73) (egg hatchability) (A) along with the percentage-point contrasts between treatments (sterile vs control and sterile males & control vs treatment for adult emergence and egg hatchability, respectively) (B). Model predictions obtained from generalized linear hierarchical models with beta and betabinomial likelihood (for adult emergence and egg hatchability, respectively) and logit link function.

For egg hatchability, the predicted mean proportion of eggs hatched was 77.1% [71.3, 82.1] for the control, 43.6 [37.7, 49.7] for the competition and 3.34% [2.39, 4.64] for the sterile males’ treatment (Figure 1A). The percentage-point contrast between the sterile males and control treatments was -73.8% [-78.9, -68.6] (p-value < 0.001; Figure 1B) while the contrast between the control and competition treatments was 33.5 [28.4, 38.6] (p-value < 0.001, Figure 1B).

#### Field trials

##### Treatment effect

Model predictions, contrasts and outputs are reported in Figure 2 and Table 3 for both crops. In radish, the predicted mean proportion of infested plants [95% confidence intervals] was 19.4% [11.7, 30.2] for the control fields and 5.6% [3.1, 10.2] for SIT-treated fields (Figure 2A). The percentage-point contrast between the treatments (control vs SIT) was -13.7 [-3.8, -23.7] (p-value = 0.00699; Figure 2B). In daikon, the predicted mean proportion of infested plants [95% confidence intervals] was 11.8% [5.2, 24.7] for the control fields and 2.1% [1.0, 4.4] for the fields with SIT (Figure 2A). The percentage-point contrast between the treatments was -9.7 [-0.8, -18.6] (p-value = 0.0327; Figure 2B).

**Table 3.** Output of statistical analyses (generalized linear hierarchical models) for the effect of Treatment (Control vs Sterile Insect Technique, SIT) and average (Avg) natural females on sticky traps per day on the proportion of radish and daikon plants infested by *Delia radicum*: estimates, corresponding 95% confidence intervals (CI), *Z* statistic and *p*-values. A betabinomial error distribution with a logit link function was used for both GLHMs. σ^2^ represents the variance of the random effects (in subscript next to value). Note that the estimates are on the logit scale.

|  | Radish |  |  |  | Daikon |  |  |  |
| --- | --- | --- | --- | --- | --- | --- | --- | --- |
| Predictors | Estimate | CI (95%) | Z | *p* | Estimate | CI (95%) | Z | *p* |
| (Intercept) | -2.82 | -3.47 – -2.16 | -8.47 | <0.001 | -3.80 | -4.58 – -3.03 | -9.67 | <0.001 |
| Treatment [control vs SIT] | 1.37 | 0.46 – 2.28 | 2.96 | **0.003** | 1.83 | 0.88 – 2.77 | 3.79 | **<0.001** |
| Avg natural females | 0.10 | -0.34 – 0.54 | 0.45 | 0.652 | 0.89 | 0.39 – 1.39 | 3.50 | **<0.001** |
| **Random Effects** |  |  |  |  |  |  |  |  |
| $\sigma^2$ | 0.34<sub>field</sub> | | | | 1.68<sub>field:year</sub> | | | |
|  |  |  |  |  | 0.16<sub>year</sub> |  |  |  |
| Observations | 220 |  |  |  | 632 |  |  |  |
| Marginal R<sup>2</sup> / Conditional R<sup>2</sup> | 0.184 / 0.304 |  |  |  | 0.502 / 1.000 |  |  |  |

**Figure 2.**
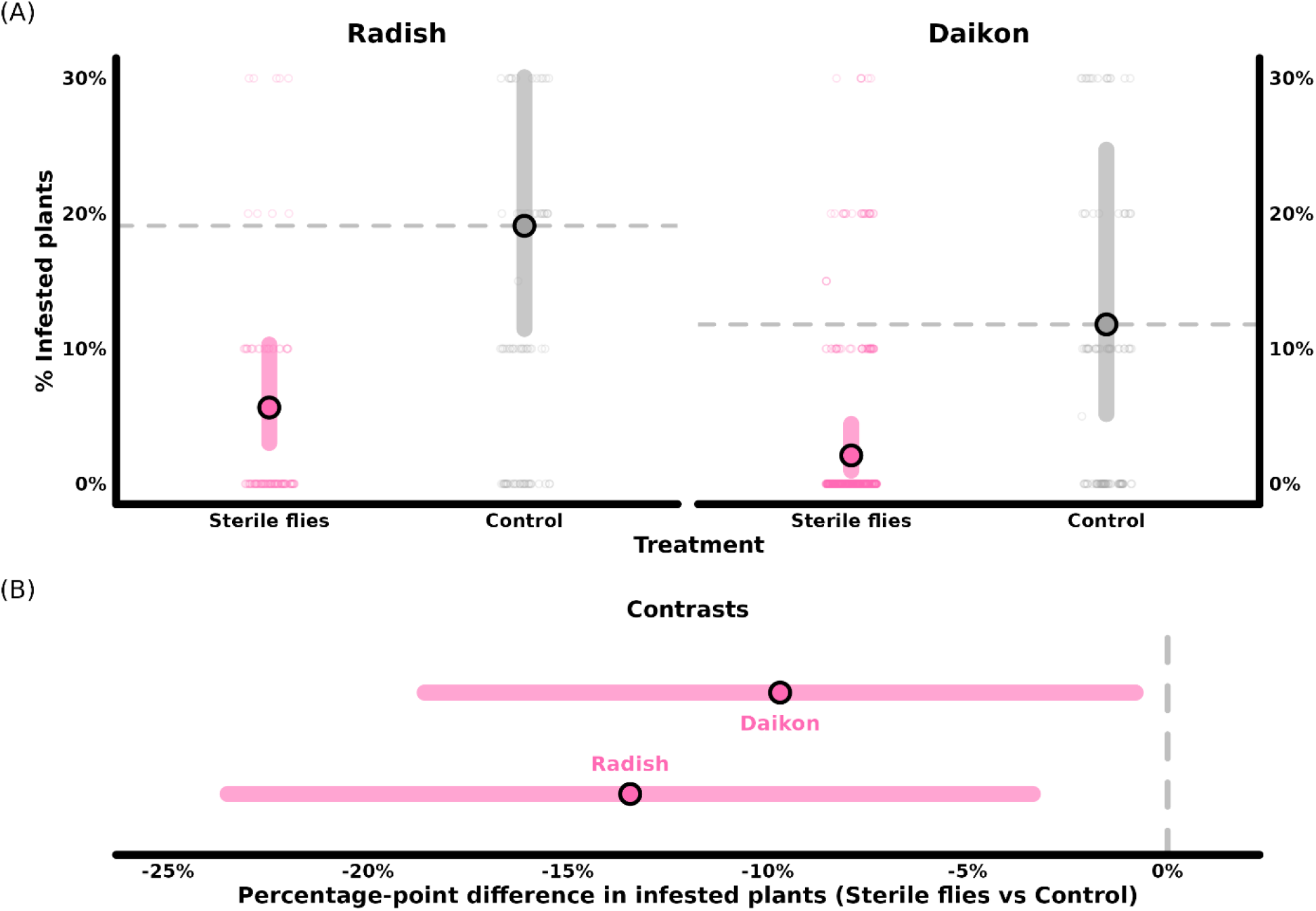
Marginal predictions of percent infested plants by D. radicum in fields with and without sterile insect technique (A) and percentage-point contrasts between treatments (control vs SIT; B) in radish and daikon fields. Model predictions obtained from generalized linear hierarchical models with betabinomial likelihood and logit link function. Total samples of 10 plants = 220 and 632 for radish and daikon, respectively. Panel (A) is cropped for aesthetic purposes. Full range of observations can be found in supplemental figure 1.

##### Control variables

The average effect (across both treatments) of the average number of natural females per sticky trap per day for the season was near null for radish whereas there was nearly 80 percentage-point increase in damage across the range of observed number of natural females in daikon (Figure 3, Table 3).

**Figure 3.**
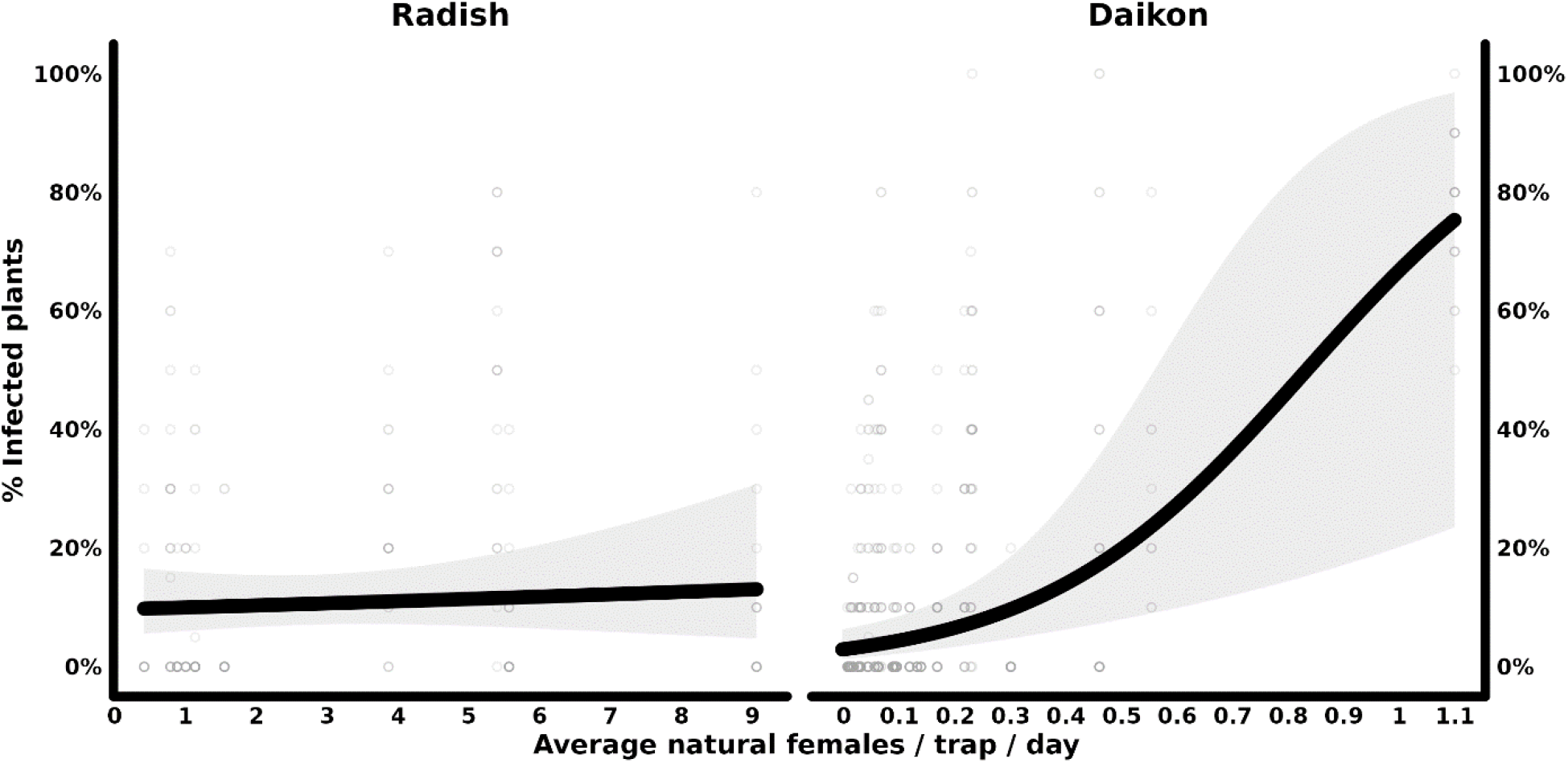
Average marginal predictions of percent infested plants by D. radicum as predicted by the average number of natural females per sticky trap per day across the growing season for radish and daikon fields. Average predictions computed for both control and SIT fields from generalized linear hierarchical models with betabinomial likelihood and logit link function. Total samples of 10 plants = 220 and 632 for radish and daikon, respectively.

## Discussion

Developing alternative pest control methods is crucial for sustainable agriculture. While the SIT has been effectively used to manage the onion maggot in Canada and abroad (Ticheler et al. 1980, Fortier et al. 2024), this study is the first to demonstrate its potential for controlling the CM and reducing it infestation rate in large-scale field trials. Our findings highlight that sterilization methods efficiently produced sterile males with minimal impact on emergence rates and mating competitiveness, ultimately leading to significant reductions in plant infestations in radish and daikon fields.

In our quality control trials, sterile individuals showed only a minor reduction (∼5%) in emergence rates compared to non-sterile. While statistically significant, this decrease has little practical impact and could be offset by increasing release numbers. Importantly, egg hatch rates were near zero when irradiated males mated with non-irradiated females, confirming the efficacy of the sterilization process. Furthermore, the 40% reduction in hatchability observed when irradiated males competed with non-irradiated males underscores their competitiveness and mating success. Indeed, since investigations of mating habits in a few *Delia* species of agricultural importance indicate that females mate only once (Martin and McEwen 1982, Spencer et al. 1997), half of them are expected to have been mated by a sterile male if their competitiveness is not affected (Fried 1971, Calkins and Parker 2005). In this case, the egg hatching rate should theoretically be intermediate between that observed for control pairs (non-irradiated) and that of females mated with a sterile male, as observed in our results. Thus, our quality control measures indicate that CM males sterilized using our methods are indeed sterile and have the potential to both compete with wild males and mate with wild females.

Following successful sterilization of CM, our SIT field trials revealed promising results. In daikon, the farm (Farm 1) had a history of high CM infestation and sterile flies have been released yearly on that farm since 2015, with very high release rates the first years (2015 and 2016), which contributed considerably to natural populations decline before the start of the trials presented here (Figure S1). Regardless of relatively low starting populations compared to radish trials, we still observed an approximately 10 percentage-point decrease in infested daikon plants in experimental fields compared to the controls. In experimental radish fields, sterile fly releases led to a 14% reduction in infestation rates, with infested plant incidence dropping to 5.6% compared to 19.4% in control fields. It is noteworthy that these radish fields had never been exposed to SIT prior to the trials and that the release rates used in the present work were relatively low since cost for growers was a major element in rate determination. As such, further releases over an increased area are expected to yield even greater reductions in damage incidence over time.

The sterile insect technique is a highly effective pest control method, particularly when integrated into area-wide pest management strategies. As demonstrated in other parts of the world, the success of SIT scales with grower adhesion, significantly reducing costs over time (Klassen and Curtis, 2005). The significant decrease in natural CM population measured on Farm 1 before the present study had resulted in gradually lower average release rates (nb flies released per ha), going from an average of 520 k flies/ha in 2015 down to 44 k in 2019 (Fortier, unpublished). This 90% decrease in release rate over five years was the same as that observed in field trials with the onion maggot *D. antiqua* in the same agroecosystem between 2011 and 2015 (Fortier et al. 2024). The strong positive response to the SIT we report here for Farm 1 is partly due to the farm’s isolated location, which limits the potential migration of *D. radicum* from neighbouring farms cultivating cruciferous crops.

An effective SIT program also requires adequate methods of monitoring natural adult populations to calibrate the number of sterile individuals released. In our field trial statistical models, we included the average number of natural females per sticky trap per day in an effort to control for differences in pest pressure between the control and sterile release fields. Interestingly, while the correlation between infestation rates and natural fly populations was evident in daikon, it was absent in radish. This discrepancy suggests that some factors may influence the relationship between natural population intensity and damage. For example, plant density is much higher in radish (550 000 per acre) than in daikon (50 000 per acre), possibly diluting the damage potential for a given population density. In addition, the geographical proximity between farms and growth duration to maturity differed greatly for the two crops. In the case of radishes, other cruciferous fields were in the vicinity of the trials, and the short production period (3 to 4 weeks) had the advantage of reducing the potential window of opportunity for damage to occur. Additional data will be necessary to determine the best proxy for predicting the risk of damage according to the density of natural populations in radish crop.

In conclusion, our study clearly demonstrates the effectiveness of SIT under commercial conditions to control *D. radicum* in radish and daikon. It offers a glimmer of hope in controlling this devastating pest in a more sustainable manner compared to conventional insecticides and could potentially be applied to other cruciferous crops like rutabaga, broccoli, cauliflower or cabbage. This approach could help reduce growers’ vulnerability in the face of declining availability of effective conventional products and rapidly evolving regulations, as well as reduce the environmental and health risks associated with the use of insecticides. Our study not only confirms the SIT’s efficacy in managing the CM in our study system but it also highlights areas for further improvements with regards to its long-term and large-scale deployment. Future efforts should focus on optimizing mass production, including the exploration of complementary molecular technologies such as RNA interference (RNAi). Additionally, developing predictive models to calibrate release timing and rates per hectare based on pest population dynamics will further enhance program efficiency. Such models will enable growers and agronomists to tailor SIT applications to specific field conditions, maximizing pest suppression while minimizing costs.

## Acknowlegements

We are grateful to the participating farms (Delfland Inc., Les Jardins A. Guérin & Fils, Les Fermes Leclair & Frères, Les Fermes du Soleil) for their trust and access to the fields. This work was funded by Agriculture and Agri-Food Canada (Canadian Agri-Science for Horticulture 3, #ASC18/19-Activity 8) and Ministère de l’Agriculture, des Pêcheries et de l’Alimentation du Québec (project 21-005-PHYD).

