## Supplemental figure 1 for "Sterile insect technique reduces cabbage maggot (Diptera: Anthomyiidae) infestation in root crucifers in Canada"

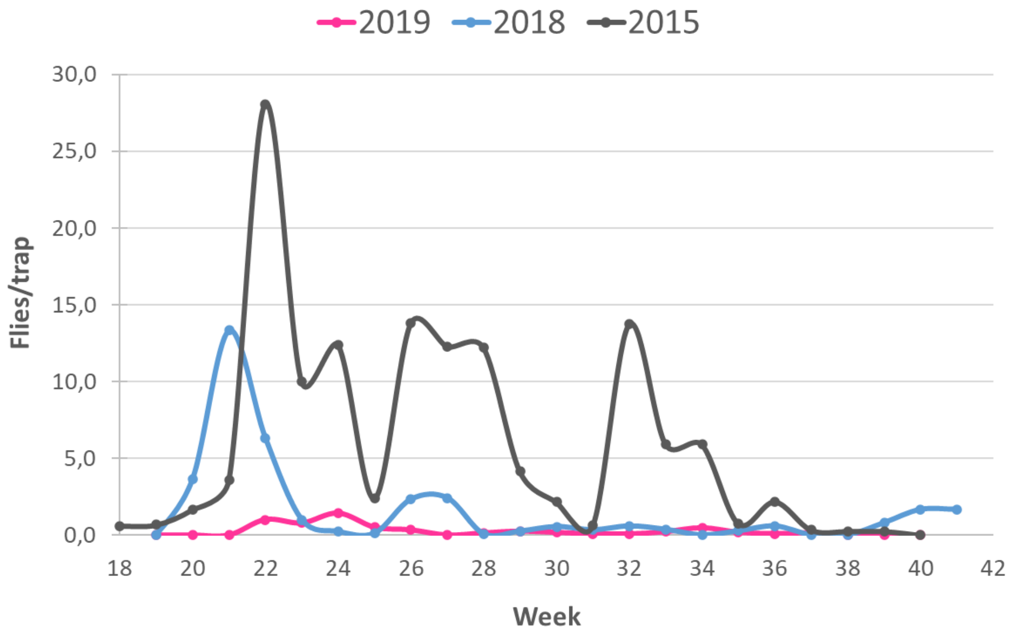


*Supplemental figure 1. Average weekly trap catches of wild D. radicum flies over seasons 2015, 2018 and 2019 in a daikon farm from Montérégie (Quebec, Canada).*
