## Supplemental figure 2 for "Sterile insect technique reduces cabbage maggot (Diptera: Anthomyiidae) infestation in root crucifers in Canada"

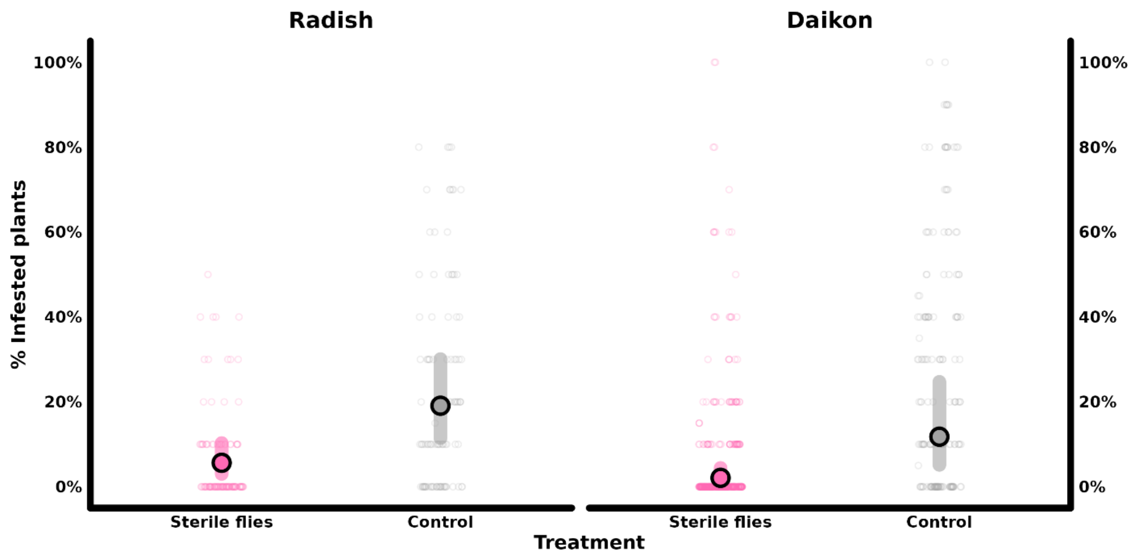


*Supplemental figure 2. Marginal predictions of percent infested plants by D. radicum in fields with and without sterile insect technique displaying entire range of observed data.*
