## Supplemental table 1 for "Sterile insect technique reduces cabbage maggot (Diptera: Anthomyiidae) infestation in root crucifers in Canada"

*Supplemental table1. Information about the area treated and release period and rate for the SIT fields monitored between 2019 and 2022 in daikon and radish crops.*

| Field | Crop | Year | Area (ha) | Release period | Nb of releases | Flies/ha (000) |
|---|---|---|---|---|---|---|
| 1 | daikon | 2019 | 10.0 | May 7 – July 9 | 10 | 75 |
| 2 | daikon | 2019 | 4.0 | July 30 – Sept. 17 | 8 | 30 |
| 3 | daikon | 2020 | 9.6 | May 5 – July 10 | 10 | 60 |
| 4 | daikon | 2020 | 6.2 | June 5 – July 24 | 8 | 35 |
| 5 | daikon | 2020 | 6.0 | June 26 – Aug. 6 | 7 | 27 |
| 6 | daikon | 2020 | 7.4 | July 24 – Sept. 17 | 9 | 25 |
| 7 | daikon | 2020 | 8.3 | Aug. 6 – Sept. 24 | 8 | 21 |
| 8 | daikon | 2020 | 1.7 | Aug. 14 – Sept. 24 | 7 | 72 |
| 9 | daikon | 2021 | 7.7 | May 3 – June 28 | 9 | 60 |
| 10 | daikon | 2021 | 2.7 | May 17 – July 5 | 8 | 50 |
| 11 | daikon | 2021 | 6.7 | May 24 – July 12 | 8 | 35 |
| 12 | daikon | 2021 | 10.5 | June 14 – Aug. 2 | 8 | 15 |
| 13 | daikon | 2021 | 10.0 | July 5 – Aug. 9 | 6 | 10 |
| 14 | daikon | 2021 | 6.7 | July 19 – Sept. 13 | 7 | 11 |
| 15 | daikon | 2021 | 4.0 | Aug. 9 – Sept. 27 | 6 | 22 |
| 16 | daikon | 2021 | 7.3 | Aug. 30 – Sept. 27 | 5 | 8 |
| 17 | daikon | 2022 | 12.0 | May 23 – July 25 | 10 | 20 |
| 18 | daikon | 2022 | 12.7 | June 20 – Aug. 22 | 10 | 11 |
| 19 | daikon | 2022 | 6.1 | July 25 – Sept. 12 | 8 | 15 |
| 20 | daikon | 2022 | 6.4 | July 18 – Sept. 6 | 8 | 17 |
| 21 | daikon | 2022 | 11.5 | Aug. 8 – Sept. 26 | 8 | 16 |
| 22 | daikon | 2022 | 4.1 | Sept. 6 – Sept. 26 | 4 | 16 |
| 1 | radish | 2020 | 11.5 | May 4 – July 6 | 9 | 50 |
| 2 | radish | 2021 | 10.0 | May 4 – June 22 | 8 | 45 |
| 3 | radish | 2021 | 6.0 | June 8 – July 20 | 7 | 70 |
| 4 | radish | 2021 | 3.0 | June 14 – July 12 | 5 | 85 |
| 5 | radish | 2022 | 4.9 | May 9 – June 6 | 5 | 50 |
